# Relapse prevention through health technology program reduces hospitalization in schizophrenia

**DOI:** 10.1101/626663

**Authors:** Philipp Homan, Nina R. Schooler, Mary F. Brunette, Armando Rotondi, Dror Ben-Zeev, Jennifer D. Gottlieb, Kim T. Mueser, Eric D. Achtyes, Susan Gingerich, Patricia Marcy, Piper Meyer-Kalos, Marta Hauser, Majnu John, Delbert G. Robinson, John M. Kane

**Affiliations:** Center for Psychiatric Neuroscience, Feinstein Institute for Medical Research, Manhasset, NY; Division of Psychiatry Research, Zucker Hillside Hospital, Northwell Health, New York, NY; Department of Psychiatry, Donald and Barbara Zucker School of Medicine at Northwell/Hofstra, Hempstead, NY; SUNY Downstate Medical School, Brooklyn, NY; Center for Technology and Behavioral Health, Dartmouth Psychiatric Research Center, Geisel School of Medicine at Dartmouth College, Lebanon, New Hampshire; Department of Critical Care Medicine, Clinical and Translational Sciences Institute, Pittsburgh, and with the Mental Illness Research, Education and Clinical Center, U.S. Department of Veterans Affairs Medical Center, Pittsburgh; Behavioral Research in Technology and Engineering (BRiTE) Center, Department of Psychiatry and Behavioral Sciences, University of Washington School of Medicine, Seattle, WA; Center for Psychiatric Rehabilitation, Boston University, Boston; Cherry Health and Pine Rest Christian Mental Health Services, Grand Rapids, MI; Division of Psychiatry and Behavioral Medicine, Michigan State University College of Human Medicine, Grand Rapids, MI; Independent consultant and trainer in Narberth, Pennsylvania; Vanguard Research Group; Minnesota Center for Chemical and Mental Health, School of Social Work, University of Minnesota, Twin Cities, St. Paul MN; Department of Mathematics, Hofstra University, Hempstead, NY

## Abstract

**Importance:** Psychiatric hospitalization is a major driver of cost in the treatment of schizophrenia. Symptom relapses are a frequent cause of hospitalization and both are primary source of burden to patients and their supporters.

**Objective:** To determine whether a novel, multicomponent, and technology-enhanced approach to relapse prevention in outpatients following a psychiatric hospitalization could reduce days spent in a hospital after discharge.

**Design:** The Improving Care and Reducing Cost (ICRC) study was a quasi-experimental clinical trial in outpatients with schizophrenia conducted between February 2013 and April 2015 at 10 different sites in the US. Data were obtained from 89 participants who received usual relapse prevention services, followed by a second cohort of 349 participants who received the technology-enhanced relapse prevention program. Both groups were followed for 6 months.

**Setting:** Outpatient setting.

**Participants:** Patients were between 18 and 60 years old; had a diagnosis of schizophrenia, schizoaffective disorder, or psychotic disorder not otherwise specified; and were currently hospitalized or had been hospitalized within the past 30 days.

**Intervention:** Patients received usual care or a technology-enhanced relapse prevention program during a 6-month period after discharge.

**Main Outcome(s) and Measure(s):** Days spent in a psychiatric hospital during 6 months after discharge.

**Results:** The study included 438 patients. Control participants (*N* = 89; 37 females) were enrolled first and received usual care for relapse prevention, and followed by 349 participants (128 females) who received technology-enhanced relapse prevention. Days of hospitalization were reduced by 4 days (Mean days: b = −4.25, 95% CI: −8.29; −0.21, *P* = 0.039) during follow-up in the intervention condition compared to control.

**Conclusions and Relevance:** The reduction in days spent in the hospital for participants in the technology-informed relapse prevention program compared to those who received usual care, and the previously reported high satisfaction and usability suggest that technology-enhanced relapse prevention is an effective and feasible way to reduce rehospitalization days among patients with schizophrenia.

**Key points:** *Question:* Can rehospitalizations in schizophrenia be prevented at reduced cost by innovative mobile technology-delivered interventions?

*Findings:* In this clinical trial with 438 patients, a technology-enhanced relapse prevention program compared to usual services reduced an average of four days of hospitalization per patient during the first 6 months following an index hospitalization.

*Meaning:* Relapse prevention through a health technology may improve care while reducing costs associated with hospitalization.

## Introduction

Technology-assisted treatment, incorporating web sites, smartphone applications, and self-paced self-management strategies, offers the promise of increasing access to and reach of potentially effective interventions in medicine. Different types of technology platforms may be appropriate for different purposes and users, and multiple interventions may be appropriate for individuals with complex, chronic conditions.^1, 2^ Here, we report outcomes for the Center for Medicaid Services-funded Improving Care and Reducing Cost (ICRC) study (Clinical Trials #NCT02364544) that used multiple technologies to prevent psychiatric hospitalization for patients with schizophrenia who had recently been discharged following a hospital admission, either to inpatient services or a day hospital.

Schizophrenia is an important disease target for chronic condition management. Psychiatric hospitalization is a major driver of cost,^3^, relapses and persistent positive symptoms impede recovery and therapeutic alliance,^4–6^, and decrease patient quality of life.^7^ For individuals with schizophrenia, poor treatment adherence, discontinuation, and service rejection are common.^8^ These characteristics drive the need for a novel multicomponent approach to patient engagement and relapse prevention:^9, 10^ The six-month “health technology program” (HTP; Supplementary Methods) included in-person, individualized relapse prevention planning and treatment that also involved treatments delivered via smartphones and computers, as well as a web-based prescriber decision support program (Fig. 1a). We hypothesized that HTP would decrease rehospitalization days during the 6-month intervention period after a recent hospitalization compared to usual care.

**Figure 1:**
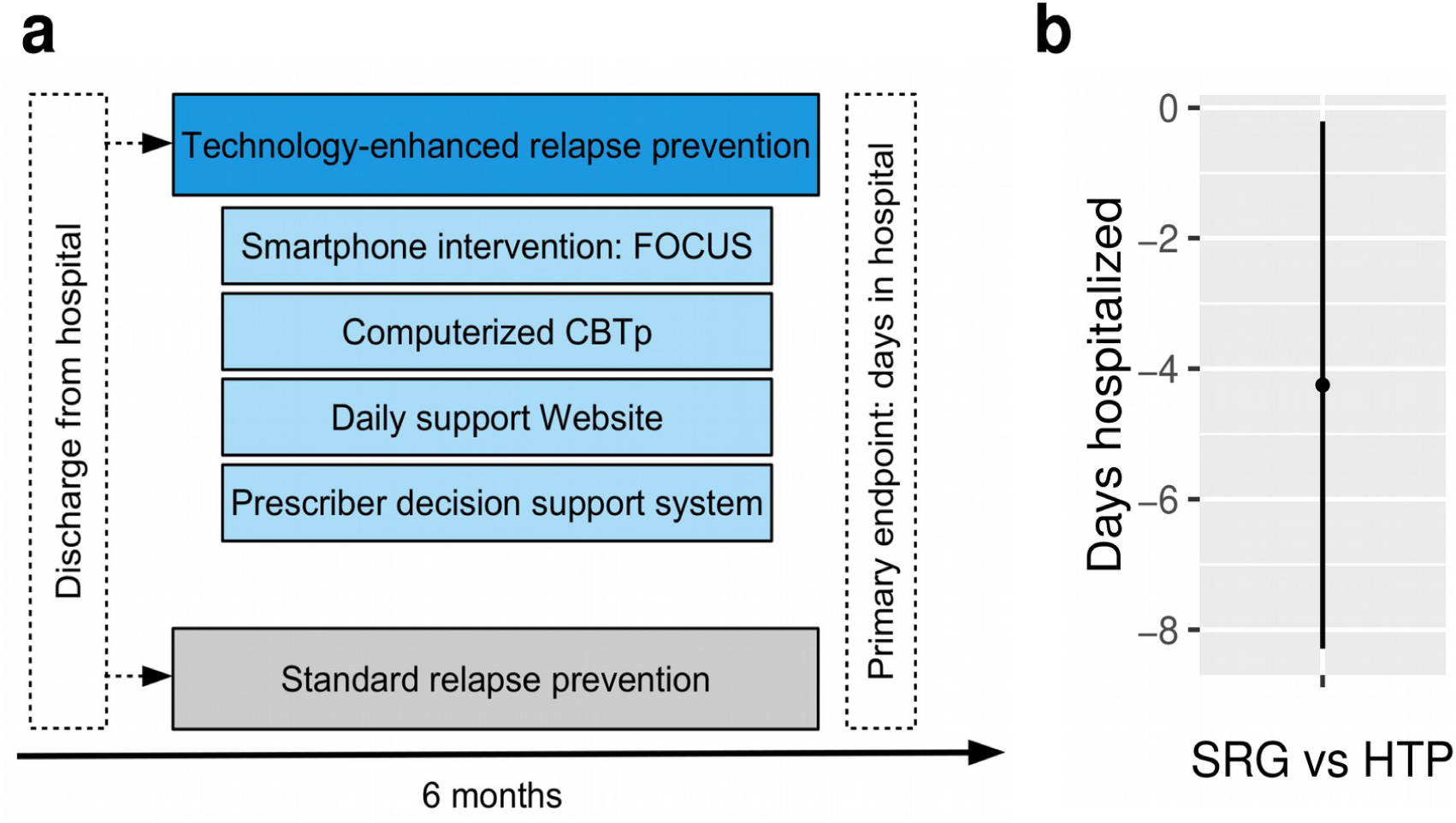
Health technology program reduces rehospitalization. **a. Study design.** After discharge from the psychiatric hospital across 10 US sites, patients were non-randomly assigned for 6 months to either the standard relapse prevention group (SRG) or the health technology program (HTP). A mental health technology coach (MHTC) guided the patient through the program, providing coaching, monitoring, and close coordination with a psychiatric prescriber. Note that HTP included (1) contacts with the MHTC to develop and maintain a relapse prevention plan; (2) an interactive smartphone illness self-management system providing coping strategy training and brief interventions to improve medication adherence, mood regulation, sleep, social functioning, and coping with auditory hallucinations; (3) a daily support web site for patients and relatives focusing on psychoeducation; (4) a web-based, self-administered, self-paced cognitive behavioral therapy education sessions for psychosis, coping with voices, and coping with paranoia; and (5) evidence-based pharmacological treatment. **b. Significant reduction in days spent in the hospital after discharge.** Compared to the SRG, the 6 month HTP significantly reduced the mean stay in psychiatric hospitals after discharge. Mean cumulative difference in days during 6 months with 95% confidence intervals is shown.

## Materials and Methods

### Study participants and design

Inclusion criteria were: age between 18 and 60 years; diagnosis of schizophrenia, schizoaffective disorder, or psychotic disorder not otherwise specified; discharge from psychiatric hospitalization within the past 30 days; having had 2 or more inpatient psychiatric admissions, ability to participate in research assessments in English; and ability to provide informed consent. Patients with a serious general medical condition were excluded.

The study was conducted at 10 sites in eight US states. Following a quasi-experimental design, up to 10 patients at each site were enrolled and assessed to form a control group starting in February 2013. After completion of control group enrollment, sites began patients enrollment in the experimental condition. Enrollment ended in April 2015. All patients provided written informed consent. The Feinstein Institutional Review Board (IRB) approved and monitored the overall study; if required by a given site, a local IRB also approved the study.

### Primary outcome

The primary outcome measure was days spent as an inpatient in a psychiatric hospital during 6 months after discharge, as assessed by a monthly patient interview.

### Statistical analysis

We used a linear mixed model with inverse probability of treatment weighting (IPTW) to estimate the treatment effect (Supplementary Material). A custom contrast comparing the difference of cumulative days spent in the hospital over the 6 month period between groups was then calculated. This contrast is reported as raw estimate (difference in days) with 95% confidence intervals. We also report balance diagnostics of the IPTW^12^ which compare baseline covariates before and after applying inverse propensity weights. We considered successful removal of confounding conditional on the IPTW by differences between covariates that were smaller than Cohen’s *d* = 0.1 for continuous covariates and smaller than Cramer’s *V* = 0.1 for categorical covariates.^12^ Faninally, we also tested for an effect of HTP on quality of life at month 6, using a IPTW-weighted linear regression and baseline quality of life as a covariate.

### Data and code availability

The data and code of this study are shared online to ensure reproducibility, doi: 10.17605/ OSF.IO/YHVCE.

## Results

Characteristics of the 89 participants assigned to the SRG group and 349 to the HTP program are presented in Table 1. Weighting the baseline covariates with inverse propensity scores resulted in a more balanced study sample,^12^ with all covariates showing effect size differences smaller than Cohen’s *d* = 0.1 and Cramer’s *V* = 0.1, respectively (Fig. S1). In addition, both groups showed high rates of study completion, with a higher rate in the HTP group that did not reach statistical significance (χ^2^ (1) = 2.9, *P* = 0.088; Table 1). Over six months and compared to the SRG group, the health technology program reduced the days of hospitalization by 4 (Mean days: b = −4.25, 95% CI: −8.29; − 0.21, *P* = 0.039; Fig. 1b). Mean estimated days of hospitalization during 6 months for the SRG group were 15.93 and 11.68 for the HTP group. Finally, using Heinrichs Carpenter Quality of Life total score at month 6 as an outcome, we found no significant effect of HTP (β = 0.02, *t* (345) = 0.43, *P* = 0.668).

**Table 1:**
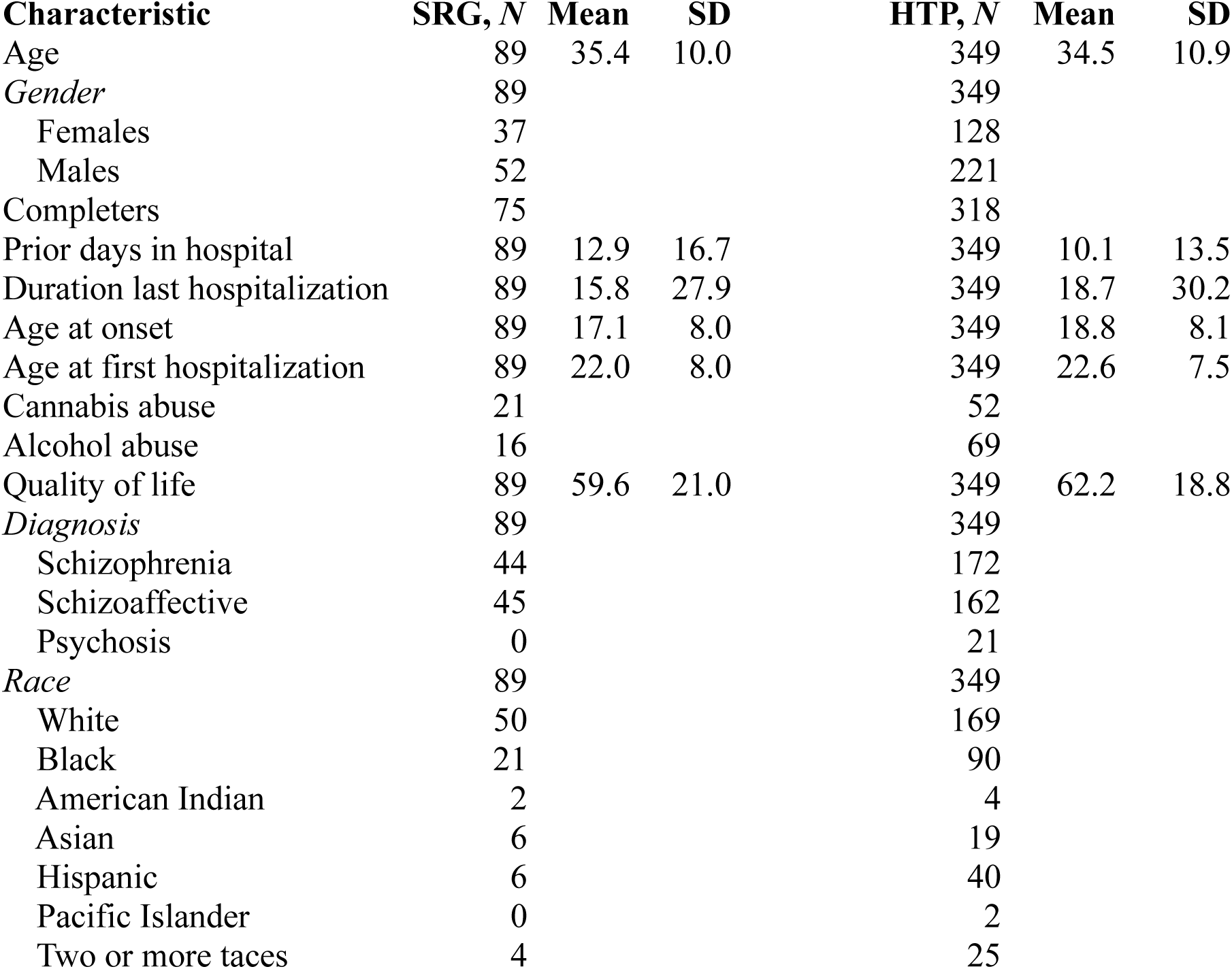
Sample characteristics of the ICRC study. *Abbreviations*: ICRC, improving care and reducing cost; SRG, standard relapse prevention group; HTP, heath technology program; Note that effect sizes for the differences between groups are shown in the Supplementary Material (Fig. S1) alongside with differences weighted for inverse propensity scores. None of these differences were significantly different between groups.

## Discussion

Recently hospitalized patients with schizophrenia who received an integrated technology-informed relapse prevention program (HTP) experienced fewer days in the hospital compared to those who received usual care in the six months following their discharge. Given the high patient burden and costs of even a single day spent in a psychiatric hospital, estimated at $1358 per day based on inflation adjusted results from a recent study^13^, our findings imply total savings in psychiatric inpatient expenditures of $5772 during the first 6 months after discharge on average. This has potentially important health-economic implications, even after additional intervention costs are taken into consideration. Technology-enhanced relapse prevention that is tailored to the individual patient may improve recovery and the ability of individuals to remain in the community, and reduce the costs for the management of schizophrenia significantly, offsetting technology access costs. In addition, the high acceptance and satisfaction reported previously^10^ suggest that these patients are willing and able to engage in a novel and technology-enhanced approach to relapse prevention.

Four features of HTP were likely to drive the beneficial impact of the intervention.^10^ First, HTP included relapse prevention planning, cognitive-behavioral symptom management strategies, family psychoeducation strategies and a computerized decision support system for medication selection, all of which are evidence-based and address common symptoms and distress. Second, in-person assistance through the MHTC may have enhanced the use of the digital tools. Third, patients’ ratings suggested that the design of the tools was helpful (e.g., easy to use for those with cognitive deficits common in schizophrenia). Finally, the HTP approach to relapse prevention was flexible, with most contacts between the MHTC and patient occurring via technologies such as mobile phones, text, and e-mail.

Although the non-randomized study design is a limitation, the high satisfaction and usability reported previously^10^ as well as low dropout suggests that HTP is a feasible and potentially effective way to increase engagement with treatment and reduce hospitalization days in schizophrenia. Thus, technology-based treatments^14^ that involve the support of trained personnel^15^ may be an efficient alternative to conventional relapse prevention approaches that are hampered by the limited availability of highly trained staff. More rigorously controlled research is needed to evaluate the effects of the HTP program, and other technology-informed programs for reducing relapses and rehospitalizations for persons with schizophrenia at risk for rehospitalization.

Much of current medical practice involves treatment of chronic conditions that impair patients in multiple domains and thereby require complex interventions. Schizophrenia, a disease often associated with motivational, cognitive and executive dysfunctions, is an example of a condition with such treatment complexities. Given the success of the HTP model for participants with schizophrenia, treatment models with multiple technologies that allow for individualization of services, which are coordinated and facilitated by a health coach, may be feasible in the management of other chronic diseases requiring complex management strategies.

## Acknowledgments

Supported by grant 1C1CMS331052-01-00 from the Centers for Medicare and Medicaid Services, Department of Health and Human Services. The contents of this report are solely the responsibility of the authors and do not necessarily represent the official views of the U.S. Department of Health and Human Services or any of its agencies.

## Supplementary Material

### Supplementary Methods

#### Health technology program

The 6-month HTP program consisted of medication treatment guided by a computer decision support system for the prescriber, a smartphone application for patients that supported medication adherence and other coping strategies, a web-based patient and family psycho-educational intervention, and web-accessed cognitive behavioral therapy for paranoia and hallucinations. A mental health technology coach provided technical support, and developed a personalized, structured, relapse prevention plan with each participant that identified individual relapse precipitants and determined which HTP components should be employed to address them. Access to the interventions was insured by providing computers and Android smartphones to all patients.

#### Inverse probability of treatment weighting

Weighting by the inverse probabilities of treatment is a method based on propensity scores,^12^ which are defined as the probability of receiving a treatment given measures of baseline covariates:

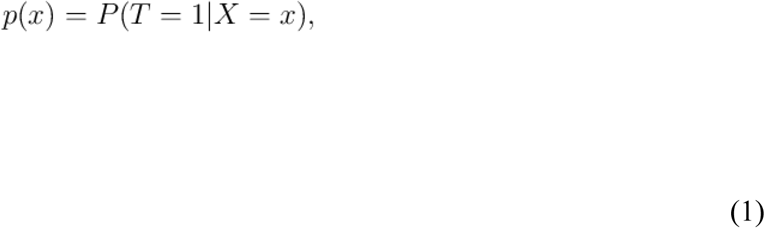

where *T* is an indicator variable that denotes the treatment received (*T*=0 for control treatment and *T*=1 for active treatment) and *x* is a vector of baseline covariates. Thus, the propensity score expresses the probability of receiving a treatment given a set of observed baseline covariates. Propensity scores are a way to address the confounding of the treatment effect that is often observed in quasi-experimental studies: treated participants will often differ systematically from untreated participants. This means that we cannot compute an unbiased estimate of the treatment effect simply by comparing outcomes between the two treatment groups:

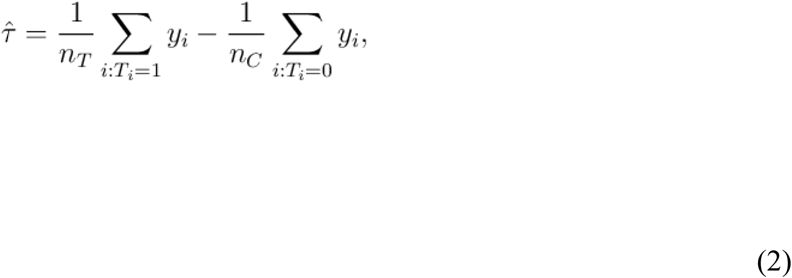

where *n*_*T*_ and *n*_*C*_ are the number of participants in the treatment and control group, respectively. When baseline confounders are present, the above treatment effect τ will not be unbiased for the true population treatment effect. Under conditional exchangability, i.e., when *Y*(1),*Y*(0)⊥*T*|*X*, we assume that when we condition on all the necessary confounders *X* that have been measured at baseline, the treatment assignment *T* is independent of the potential outcome *Y*. Thus, by conditioning on propensity scores, it is possible to balance the measured baseline covariates so that the treatment and control groups are similar with respect to these covariates.

Inverse probability weights are one way of applying propensity scores. They are derived by taking the inverse of the propensity score:

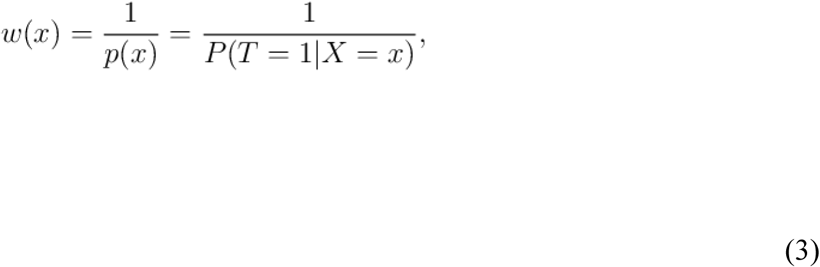

for treated individuals and

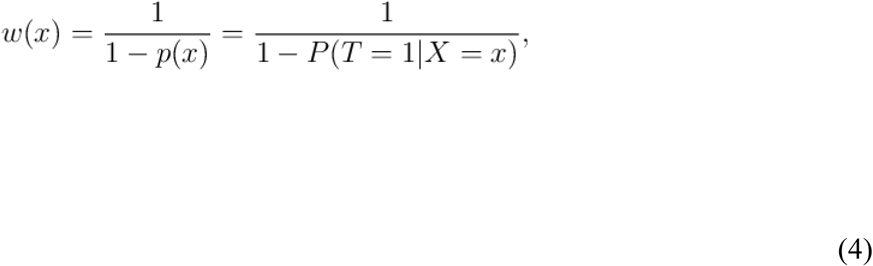

for untreated individuals. They create a synthetic sample in which the distribution of measured baseline covariates is independent of treatment assignment. To avoid that individuals with a propensity score close to 0 (i.e., those very unlikely to be treated) wound end up with overly large weights (which would make the weighted estimator unstable) we used stabilized IPTW. For treated individuals, the stabilized weights are calculated as follows:

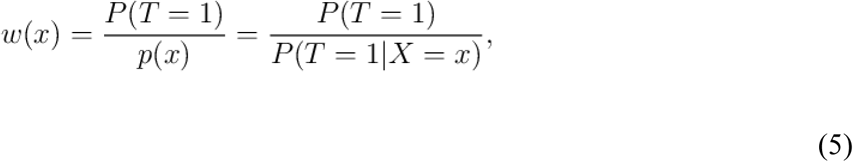

wheres for untreated subjects, they are given by

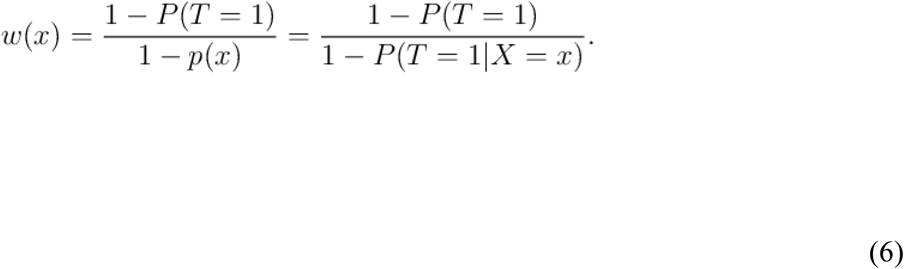

We then used these stabilized weights to weight each participants outcome, the number of days spent in a psychiatric hospital, which was the primary outcome measure, and used treatment (SRG vs. HTP) and number of visits as predictors. We also included predictors for all the covariates that were used to calculate the propensity scores, including age, gender, race, diagnosis, socioeconomic status, study site, quality of life at baseline, cannabis and alcohol abuse, age at illness onset, age at first hospitalization, number of prior hospitalizations, length of last hospitalization, and days in medical treatment.

## Supplementary Figures

**Figure 1:**
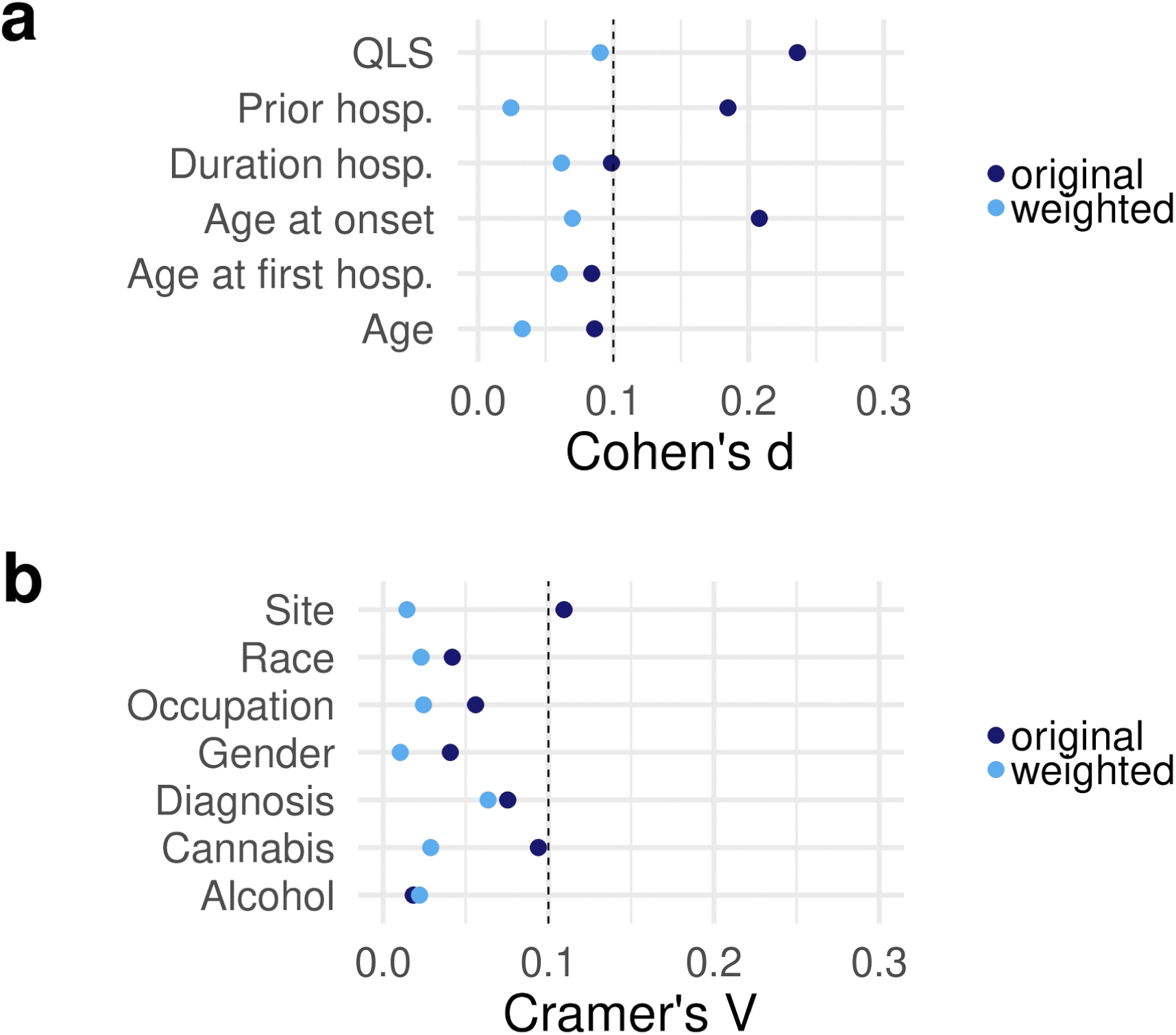
Balance diagnostics confirm removal of confounding. a, b. Continuous and categorical covariates were more similar after applying inverse propensity weights. This is indicated by weighted values being smaller than the cutoffs (indicated with dashed lines). QLS, quality of life scale; hosp., hospitalization;

